# D-Serine inhibits non-ionotropic NMDA receptor signaling

**DOI:** 10.1101/2024.05.29.596266

**Authors:** Eden V. Barragan, Margarita Anisimova, Vishnu Vijayakumar, Azariah C. Coblentz, Deborah K. Park, Raghava Jagadeesh Salaka, Atheer F.K. Nisan, Samuel Petshow, Kim Dore, Karen Zito, John A. Gray

**Author notes:** Co-corresponding Authors: John A. Gray, MD PhD; Karen Zito, PhD. Conflicts of Interest:* The authors declare no competing financial interests.

## Abstract

NMDA-type glutamate receptors (NMDARs) are widely recognized as master regulators of synaptic plasticity, most notably for driving long-term changes in synapse size and strength that support learning. NMDARs are unique among neurotransmitter receptors in that they require binding of both neurotransmitter (glutamate) and co-agonist (e.g. d-serine) to open the receptor channel, which leads to the influx of calcium ions that drive synaptic plasticity. Over the past decade, evidence has accumulated that NMDARs also support synaptic plasticity via ion flux-independent (non-ionotropic) signaling upon the binding of glutamate in the absence of co-agonist, although conflicting results have led to significant controversy. Here, we hypothesized that a major source of contradictory results can be attributed to variable occupancy of the co-agonist binding site under different experimental conditions. To test this hypothesis, we manipulated co-agonist availability in acute hippocampal slices from mice of both sexes. We found that enzymatic scavenging of endogenous co-agonists enhanced the magnitude of LTD induced by non-ionotropic NMDAR signaling in the presence of the NMDAR pore blocker, MK801. Conversely, a saturating concentration of d-serine completely inhibited both LTD and spine shrinkage induced by glutamate binding in the presence of MK801. Using a FRET-based assay in cultured neurons, we further found that d-serine completely blocked NMDA-induced conformational movements of the GluN1 cytoplasmic domains in the presence of MK801. Our results support a model in which d-serine inhibits ion flux-independent NMDAR signaling and plasticity, and thus d-serine availability could serve to modulate NMDAR signaling even when the NMDAR is blocked by magnesium.

**Significance Statement:** NMDARs are glutamate-gated cation channels that are key regulators of neurodevelopment and synaptic plasticity and unique in their requirement for binding of a co-agonist (e.g. d-serine) in order for the channel to open. NMDARs have been found to drive synaptic plasticity via non-ionotropic (ion flux-independent) signaling upon the binding of glutamate in the absence of co-agonist, though conflicting results have led to controversy. Here, we found that d-serine inhibits non-ionotropic NMDAR-mediated LTD and LTD-associated spine shrinkage. Thus, a major source of the contradictory findings might be attributed to experimental variability in d-serine availability. In addition, the developmental regulation of d-serine levels suggests a role for non-ionotropic NMDAR plasticity during critical periods of plasticity.

## Introduction

*N*-methyl-D-aspartate receptors (NMDARs) are glutamate-gated ion channels that play crucial roles in neurodevelopment and synaptic plasticity; even subtle changes in NMDAR functioning can have wide-ranging developmental and cognitive effects. NMDARs are unique among neurotransmitter receptors in that, in addition to binding of the neurotransmitter (glutamate), they require co-agonist (e.g. d-serine) binding in order to open the receptor channel pore (Hansen et al., 2018). Over the past decade, there has been growing recognition that NMDARs (Dore et al., 2016; Gray et al., 2016; Park et al., 2022a), along with other ligand-gated ion channels (Valbuena and Lerma, 2016), support agonist-mediated intracellular signaling independent of ion flux through the channels. Indeed, ion flux-independent (or non-ionotropic) forms of NMDAR-mediated synaptic plasticity have been described, including long-term depression (LTD) and dendritic spine shrinkage (Mayford et al., 1995; Nabavi et al., 2013; Stein et al., 2015; Carter and Jahr, 2016; Wong and Gray, 2018; Stein et al., 2020; Dore and Malinow, 2021; Stein et al., 2021). In these studies, NMDAR-dependent LTD and/or dendritic spine shrinkage were observed in response to glutamate binding, even when ion flux through the receptor was blocked with the uncompetitive channel blocker MK801 or with a competitive antagonist of the co-agonist site.

Despite the growing evidence supporting a role for ion flux-independent NMDAR signaling in synaptic plasticity, there remains controversy regarding the phenomenon. Notably, several studies have reported conflicting results, particularly with the use of MK801, which has been shown to inhibit LTD (Sanderson et al., 2012; Babiec et al., 2014; Coultrap et al., 2014; Sanderson et al., 2016). Intriguingly, we noticed that, in our hands, LTD induced in the presence of the pore blocker MK801 was smaller in magnitude and had more experiment-to-experiment variation as compared to LTD induced during co-agonist site inhibition. Importantly, while open-channel blockers like MK801 effectively block ion flux through the NMDAR channel, they do not significantly alter the affinity or occupancy of the co-agonist binding site (Huettner and Bean, 1988; MacDonald et al., 1991; Blanpied et al., 1997; Bolshakov et al., 2003). Thus, we hypothesized that co-agonist occupancy might modulate glutamate-induced ion flux-independent NMDAR-mediated plasticity.

Here we show that competitive antagonism of the NMDAR co-agonist site or enzymatic scavenging of endogenous co-agonists both increases the magnitude and reduces the inter-experiment variability of LTD induced in the presence of MK801. In addition, a saturating concentration of d-serine completely inhibits ion flux-independent NMDAR-mediated LTD and LTD-associated spine shrinkage and NMDA-induced conformational movements of the GluN1 cytoplasmic domains. Our results demonstrate that, surprisingly, d-serine blocks non-ionotropic NMDAR-mediated plasticity. These results suggest that experimental differences in d-serine availability could contribute to the inconsistencies observed when using MK801 to study non-ionotropic LTD. Furthermore, d-serine availability could serve to modulate NMDAR downstream signaling, even at rest when the NMDAR is blocked by magnesium.

## Materials and Methods

### Animals

Wild-type C57BL/6J mice (#000664 Jax) of both sexes were group housed in polycarbonate cages and maintained on a 12h light/dark cycle at a constant temperature of 24 ± 1°C. For two-photon imaging and uncaging experiments we used GFP-M mice (Feng et al., 2000) in a C57BL/6J background to obtain sparsely GFP-labelled pyramidal neurons in the CA1 area of the hippocampus. Serine racemase knockout (SRKO; Basu et al., 2009) and GFP-M mice in a C57BL/6J background were crossed to generate serine racemase knockout and wild-type littermates with sparsely GFP-labelled CA1 pyramidal neurons (Park et al., 2022b). Animals were given access to food and water *ad libitum*. All experiments were carried out in accordance with the National Institutes of Health guidelines and were approved by the UC Davis Institutional Animal Care and Use Committee (IACUC).

### Enzymes

Purified recombinant enzymes, *Escherichia coli* d-serine deaminase (DsdA; EC 4.3.1.18) and *Bacillus subtilis* glycine oxidase (GO; EC 1.4.3.19) were the generous gift of Herman Wolosker (Technion Institute, Israel). DsdA is a bacterial enzyme that has significant advantages over the more commonly used D-amino acid oxidase (DAAO). DsdA is at least three orders of magnitude more efficient than DAAO in destroying d-serine (Shleper et al., 2005), has a significantly higher apparent affinity for d-serine (Km = 0.1 mM) compared to DAAO (Km = 50 mM) (Shleper et al., 2005), does not produce hydrogen peroxide as a byproduct (Lu et al., 2012; Matlashov et al., 2014), and is highly specific to d-serine (Dupourque et al., 1966) thus eliminating possible off-target effects on other D-amino acid substrates. In addition, DsdA efficiently degrades d-serine in organotypic slices (Shleper et al., 2005), neuronal cultures (Kartvelishvily et al., 2006), and retina preparations (Gustafson et al., 2007). DsdA was expressed and purified as described previously (Shleper et al., 2005; Kartvelishvily et al., 2006), concentrated to 36.3 mg/mL in 10 mM Tris-HCl pH 8.5, and frozen at -70°C. DsdA was used at a final concentration of 5 µg/mL. The H244K mutant of GO, which has an 8-fold higher specific activity for glycine compared with wild-type (Rosini et al., 2014) was purified as described previously (Job et al., 2002; Settembre et al., 2003) and concentrated to 60 mg/mL (∼2 U/mg) in 10 mM Na-pyrophosphate pH 8.5 and 10% glycerol, and frozen at -70°C. GO was used at a final concentration of ∼0.12 U/mL. For enzymes experiments, slices were pre-incubated with both GO and DsdA in artificial cerebrospinal fluid (ACSF) for at least 60 min to optimally degrade endogenous co-agonists, then continuously perfused during recordings in a reduced-volume recirculating perfusion system. For controls, enzymes were heat inactivated at 65°C for 10 min.

### Electrophysiology

P14-P18 mice were anesthetized with isoflurane and decapitated. Brains were rapidly removed and placed in ice-cold sucrose cutting buffer, containing the following (in mM): 210 sucrose, 25 NaHCO_3_, 2.5 KCl, 1.25 NaH_2_PO_4_, 7 glucose, 7 MgCl_2_, and 0.5 CaCl_2_. Acute transverse 400 µm slices (300 µM for whole-cell recordings) were made by dissecting the hippocampus out of each hemisphere and mounting on agar. Slices were cut on a Leica VT1200 vibratome in ice-cold sucrose cutting buffer, then recovered for at least 1 hr in 32°C ACSF containing (in mM): 119 NaCl, 26.2 NaHCO_3_, 11 glucose, 2.5 KCl, 1 NaH_2_PO_4_, 2.5 CaCl_2_, and 1.3 MgSO_4_. Slices were stored submerged in room temperature ACSF for up to 5h, before they were transferred to a submersion chamber on an upright Olympus microscope, perfused with room temperature ACSF containing picrotoxin (0.1 mM), and saturated with 95% O_2_/5% CO_2_.

### Extracellular Field Electrophysiology

For extracellular field EPSP (fEPSP) recordings, a bipolar tungsten stimulating electrode (MicroProbes) was placed in stratum radiatum of the CA1 region and used to activate Schaffer collateral (SC)-CA1 synapses. Evoked field excitatory postsynaptic potentials (fEPSPs) were recorded in stratum radiatum using borosilicate pipettes (Sutter Instruments) filled with ACSF (resistance ranged from 5–10 MΩ) at a basal stimulation rate of 0.05 Hz). At the start of each experiment, the maximal fEPSP amplitude was determined and the intensity of presynaptic fiber stimulation was adjusted to evoke fEPSPs with an amplitude of ∼50% of the maximal amplitude. After obtaining a stable 10 min baseline, slices were stimulated using either a standard low-frequency LTD induction protocol (900 stimuli at 1 Hz), a neutral protocol (900 stimuli at 10 Hz), or a high-frequency LTP induction protocol (300 stimuli at 50 Hz) (Dudek and Bear, 1992). The mean slope of excitatory postsynaptic potentials (EPSPs) of the final 10 minutes of each recording (normalized to baseline) was used for statistical comparisons. Slices were pre-incubated for at least 60 min in 100 µM (+)-MK801 (Tocris) before recordings. MK801, 10 µM L-689,560 (L689; Tocris) and 50 µM D-AP5 (HelloBio) were bath applied throughout the entirety of the recording. 10 µM d-serine (Tocris) was removed after the induction stimulus. NMDAR-meditated fEPSPs were recorded in low Mg^2+^ (0.2 mM) ACSF following application of 10 µM NBQX to block AMPA receptors. All recordings were collected with a Multiclamp 700B amplifier (Molecular Devices). Analysis was performed with the Clampex 11.2 software suite (Molecular Devices) and GraphPad Prism 10.0 software.

### Whole-Cell Electrophysiology

CA1 neurons were visualized by infrared differential interference contrast microscopy. Cells were patched with 3-5MΩ borosilicate pipettes filled with intracellular solution, containing (in mM) 135 cesium methanesulfonate, 8 NaCl, 10 HEPES, 0.3 Na-GTP, 4 Mg-ATP, 0.3 EGTA, and 5 QX-314 (Sigma).

Excitatory postsynaptic currents (EPSCs) were evoked by electrical stimulation of Schaffer collaterals with a bipolar tungsten electrode (MicroProbes). NMDAR-EPSCs were measured at -40 mV in the presence of 10 µM NBQX. Series resistance was monitored and not compensated, and cells were discarded if series resistance varied more than 25%. All recordings were obtained with a Multiclamp 700B amplifier, filtered at 2 kHz, digitized at 10 Hz. Analysis was performed with the Clampex 11.2 software suite and GraphPad Prism 10.0.

### Two-photon imaging

Acute hippocampal slices were prepared from P16-P21 GFP-M mice (Feng et al., 2000) or SRKO (Basu et al., 2009) crossed with GFP-M of both sexes in C57BL/6J background. Coronal 300 μm slices (400 μm for SRKO experiments) were cut (Leica VT100S vibratome) in cold choline chloride dissection solution containing (in mM): 110 choline chloride, 2.5 KCl, 25 NaHCO_3_, 0.5 CaCl_2_, 7 MgCl_2_, 1.3 NaH_2_PO_4_, 11.6 sodium ascorbate, 3.1 sodium pyruvate, and 25 glucose, saturated with 95% O_2_/5% CO_2_. Slices were recovered at 30°C for 30-45 min then at room temperature (RT) for an additional 30-45 min, in oxygenated ACSF containing (in mM): 127 NaCl, 25 NaHCO_3_, 1.25 NaH_2_PO_4_, 2.5 KCl, 25 glucose, 2 CaCl_2_, and 1 MgCl_2_. GFP-expressing CA1 pyramidal neurons (30-60 µm depth) were imaged using a custom two-photon microscope (Woods et al., 2011). For each neuron, image stacks (512 x 512 pixels; 0.02 μm per pixel; 1-μm z-steps) were collected from one segment of secondary or tertiary basal dendrites at 5 min intervals in recirculating ACSF consisting of (in mM): 127 NaCl, 25 NaHCO_3_, 1.2 NaH_2_PO_4_, 2.5 KCl, 25 D-glucose, and 1.5 or 2 Ca^2+^, 0 or 1 Mg^2+^(as specified in text), aerated with 95%O_2_/5%CO_2_, ∼310 mOsm, pH 7.2) at 27-30°C with 2.5 mM MNI-glutamate (Tocris) and 1 μM TTX (Tocris). Slices were pre-incubated with pharmacological reagents prior to imaging: 1 h for enzymes (GO and DsdA, described above), heat-inactivated enzymes, MK-801 (100 μM), or Mg^2+^ (1 mM); 10-15 min for d-serine (10 µM), or NBQX (50 µM). All remained for the entire imaging experiment, except d-serine (10 µM) which was removed immediately after uncaging. Estimated spine volume was measured from background-subtracted green fluorescence using the integrated pixel intensity of a boxed region surrounding the spine head, as described (Woods et al., 2011). Representative images are maximum projections of three-dimensional image stacks after applying a median filter (3 × 3) to raw image data.

### Glutamate uncaging

Uncaging of MNI-glutamate (2.5 mM) occurred directly after two baseline images using a parked beam (720 nm, 9-12 mW at the sample) at a point ∼0.5-1 μm from the spine head. Subthreshold high-frequency glutamate uncaging (sub-HFU) consisted of 60 pulses of 2 ms at 0.5 Hz in 1.5 mM Ca^2+^.

Low-frequency glutamate uncaging (LFU) consisted of 90 pulses of 0.2 ms duration at 0.1 Hz in 2 mM Ca^2+^ and 0 mM Mg^2+^. High frequency glutamate uncaging (HFU) consisted of 60 pulses of 2 ms at 0.5 Hz in 2 mM Ca^2+^ and 0 or 1 mM Mg^2+^ (as specified in text).

### Fluorescence lifetime imaging of intracellular NMDAR conformation

C57BL/6J mouse pups (#000664 Jax) were used to prepare primary hippocampal neurons according to previously described protocols (Dore et al., 2015). Neurons were transfected at DIV 7-10 with ∼2 μg of total DNA (GluN1-GFP, GluN2B and GluN1-mCherry) and 4 μL of Lipofectamine 2000 was used per well (18mm coverslips, Neuvitro). The fluorescence lifetime of GluN1-GFP, the FRET donor, is highly sensitive to the proximity of GluN1-mCherry, the FRET acceptor; fluorescence lifetime imaging microscopy (FLIM) can be used measure Förster resonance energy transfer (FRET) in a precise and quantitative manner (Yasuda, 2006; Dore et al., 2015). Prior to imaging, 14-18 DIV neurons were incubated in 100 μM of MK801 for 1 h. Imaging was done in a HBSS based solution containing: 0.87x HBSS, 5mM HEPES, 1mM Glucose, 2.5mM MgCl_2_, 0.5mM CaCl_2_. MK801 was also added to the HBSS imaging solution along with 10 μM d-serine when specified. After 1-3 suitable neurons were found for imaging, a baseline image was recorded. The perfusion was then switched to a HBSS imaging solution containing 25 μM NMDA (Tocris) and neurons were imaged a second time. Each neuron was imaged only twice, and drugs conditions were interleaved. Results are pooled from at least two different animal preparations. FLIM was performed on a SliceScope two-photon microscope (Scientifica) as previously described (Dore et al., 2015). Briefly, a Chameleon Ultra II IR laser (Coherent) tuned at 930 nm was used for GFP excitation (power adjusted to 3mW after the microscope objective (LUMPLFLN 60XW, NA = 1.0, Olympus)). Fluorescence emission was detected with a hybrid PMT detector (HPM-100-40, Becker and Hickl, Germany) and synchronized by a TCSPC module (SPC-150, Becker and Hickl); all imaging parameters were kept constant for all acquired images. FLIM images were processed with SPCImage (Becker and Hickl) and further analyzed blind to condition with a custom MATLAB script (Dore et al., 2015).

### Experimental rigor and statistical analysis

A minimum of three mice were used per group. With the exception of the enzyme electrophysiology experiments, controls and drug treatments were interleaved each day. Statistical comparisons were made with unpaired *t* tests or with one-way or two-way ANOVAs with Bonferroni’s post-hoc multiple comparisons test as specified and appropriate, using GraphPad Prism 10.1 with *p* < 0.05 considered significant.

## Results

### Decreasing co-agonist site occupancy reduces the variability and enhances the magnitude of ion flux-independent NMDAR-LTD

In order to test our hypothesis that occupancy of the co-agonist site could influence the magnitude of ion flux-independent NMDAR-mediated plasticity, we first took a closer quantitative look at a comparison between LTD in the presence of the NMDAR pore blocker, MK801, and LTD in the presence of the NMDAR co-agonist site competitive antagonist, L-689,560 (L689). Indeed, we observed that LTD in the presence of MK801 was of lower magnitude than LTD observed in the presence of L689 (**Fig. 1A**; MK801: 86.3±3.6%, *n*=19; +L689: 75.1±1.8%, *n*=16; *p*=0.022, one-way ANOVA with Bonferroni post-hoc multiple comparisons test). In addition, the experiment-to-experiment variability was higher with MK801 alone compared to MK801 with L689 (**Fig. 1B**). Similarly, applying both inhibitors simultaneously, we again found that addition of the co-agonist site antagonist L689 results in a modest but significant increase in the magnitude of LTD compared to MK801 alone (**Fig. 1C**; MK801: 86.0±4.2%, *n*=16; +L689: 73.6±2.4%, *n*=9; *p*=0.0331, one-way ANOVA with Bonferroni post-hoc multiple comparisons test), and a reduction in experiment-to-experiment variability (**Fig. 1D**). Importantly, 100 µM MK801 completely inhibits NMDAR-meditated fEPSPs (**Fig. 1E-F**; ACSF: 12.7±1.5% of baseline, *n*=7, *p*=0.0002; MK801: 0.36±1.3%, *n*=6, *p*=0.7925, one sample *t* test compared to a hypothetical mean of zero).

**Figure 1.**
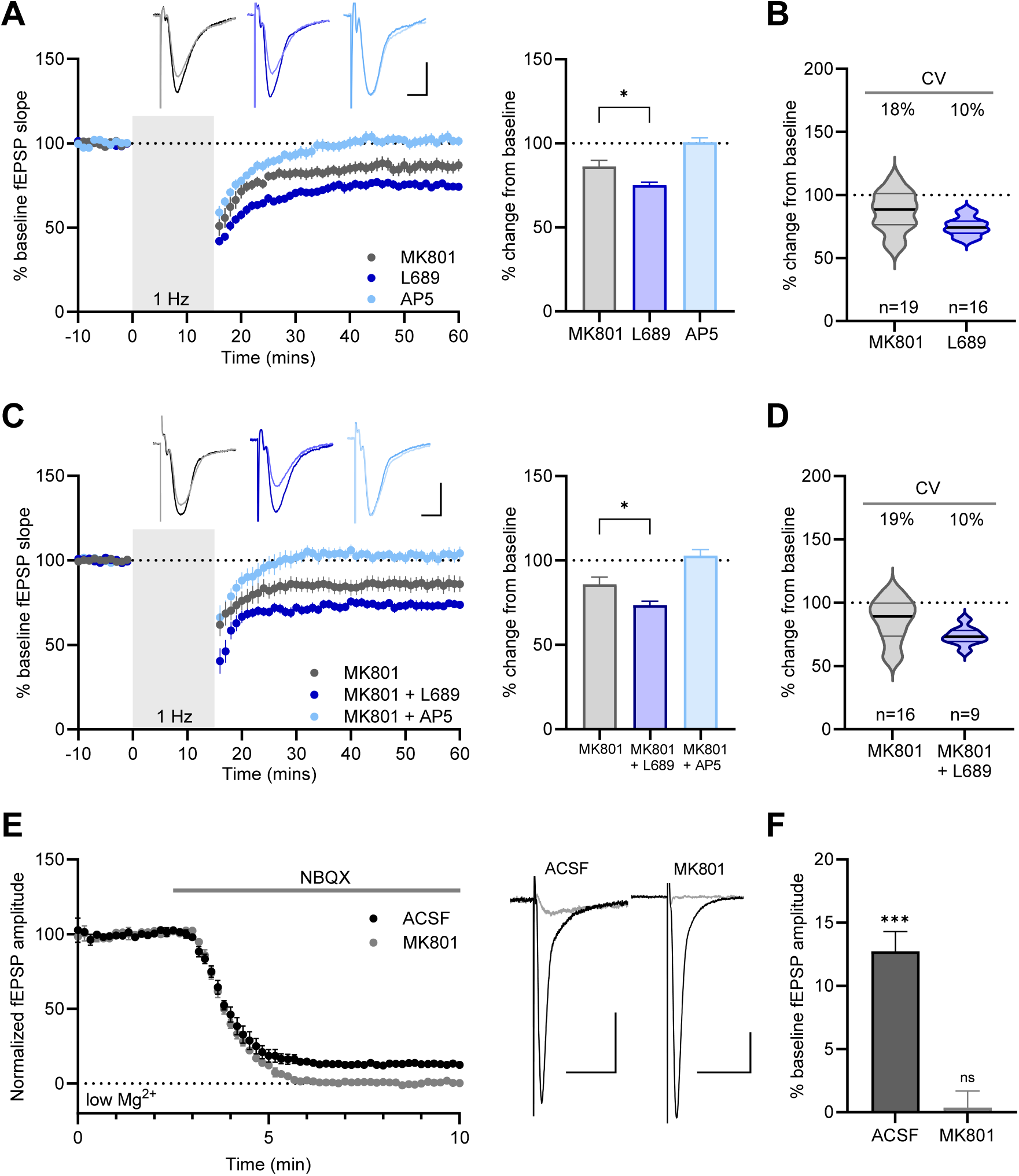
Co-agonist site antagonism increases the magnitude and reduces the variance of ion flux-independent NMDAR-mediated LTD. **A**, *Left:* averaged plasticity experiments using a 1 Hz, 900 pulse LTD induction protocol in the presence of 100 µM MK801 (grey circles; *n*=19), 10 µM L689 (dark blue circles; *n*=16) or 50 µM AP5 (light blue circles; *n*=12). *Right:* Compared with MK801, L689 resulted in an increased magnitude of LTD (unpaired *t* test). **B,** Coefficient of variation (CV) of the experiments in A demonstrating the reduction in variation with L689 compared to MK801. **C**, *Left:* averaged plasticity experiments using a 1 Hz, 900 pulse LTD induction protocol in the presence of 100 µM MK801 alone (grey circles; *n*=16), or with the addition of either 10 µM L689 (dark blue circles; *n*=9) or 50 µM AP5 (light blue circles; *n*=9). *Right:* Compared with MK801 alone, the addition of L689 resulted in an increased magnitude of LTD (unpaired *t* test). **D,** Coefficient of variation (CV) of the experiments in A demonstrating the reduction in variation with L689 compared to MK801 alone. **E,** *Left:* averaged NMDAR fEPSPs recorded in 0.2 mM Mg^2+^ normalized to baseline fEPSP amplitude prior to blocking AMPA receptors with 10 µM NBQX in slices pre-incubated with ACSF (black circles; *n*=7) or 100 µM MK801 (grey circles; *n*=6). *Right:* sample traces before (black) and after (grey) NBQX. **F,** summary data of E, MK801 pre-incubation completely blocks NMDAR fEPSPs (one sample *t* test compared to 0). Scale bars for all sample traces are 0.5 mV, 10 msec. All data represented as mean ± SEM. \**p*<0.05, \*\*\**p*<0.001, ^ns^*p*≥0.05.

The higher variability and lower magnitude of ion flux-independent NMDAR-mediated LTD with MK801 suggests that some other factor is modulating the success rate of ion flux-independent LTD that is eliminated in the presence of the co-agonist site inhibitor L689, such as variable occupancy of the co-agonist binding site in the presence of MK801 (Huettner and Bean, 1988; MacDonald et al., 1991; Blanpied et al., 1997; Bolshakov et al., 2003). However, an alternative explanation could be that co-agonist site competitive antagonists may be stabilizing conformational states distinct from an unoccupied site. Thus, we performed a similar experiment, using enzymatic scavenging of endogenous co-agonists from brain slices (Mothet et al., 2000; Panatier et al., 2006; Papouin et al., 2012; Li et al., 2013; Le Bail et al., 2015; Meunier et al., 2016) with recombinant d-serine deaminase (DsdA) and glycine oxidase (GO). Hippocampal slices were pre-incubated for at least 60 min with both DsdA and GO, then continuously perfused with enzymes during recordings using a recirculating perfusion system. Like L689, enzymatic scavenging of endogenous co-agonists results in a significant increase in the magnitude of non-ionotropic LTD compared to MK801 alone (**Fig. 2A**; MK801: 87.7±5.1%, *n*=11; +enzymes: 73.1±3.2%, *n*=8; *p*=0.0497, one-way ANOVA with Bonferroni post-hoc multiple comparisons test) and similarly reduced the experiment-to-experiment variability (**Fig. 2B**).

**Figure 2.**
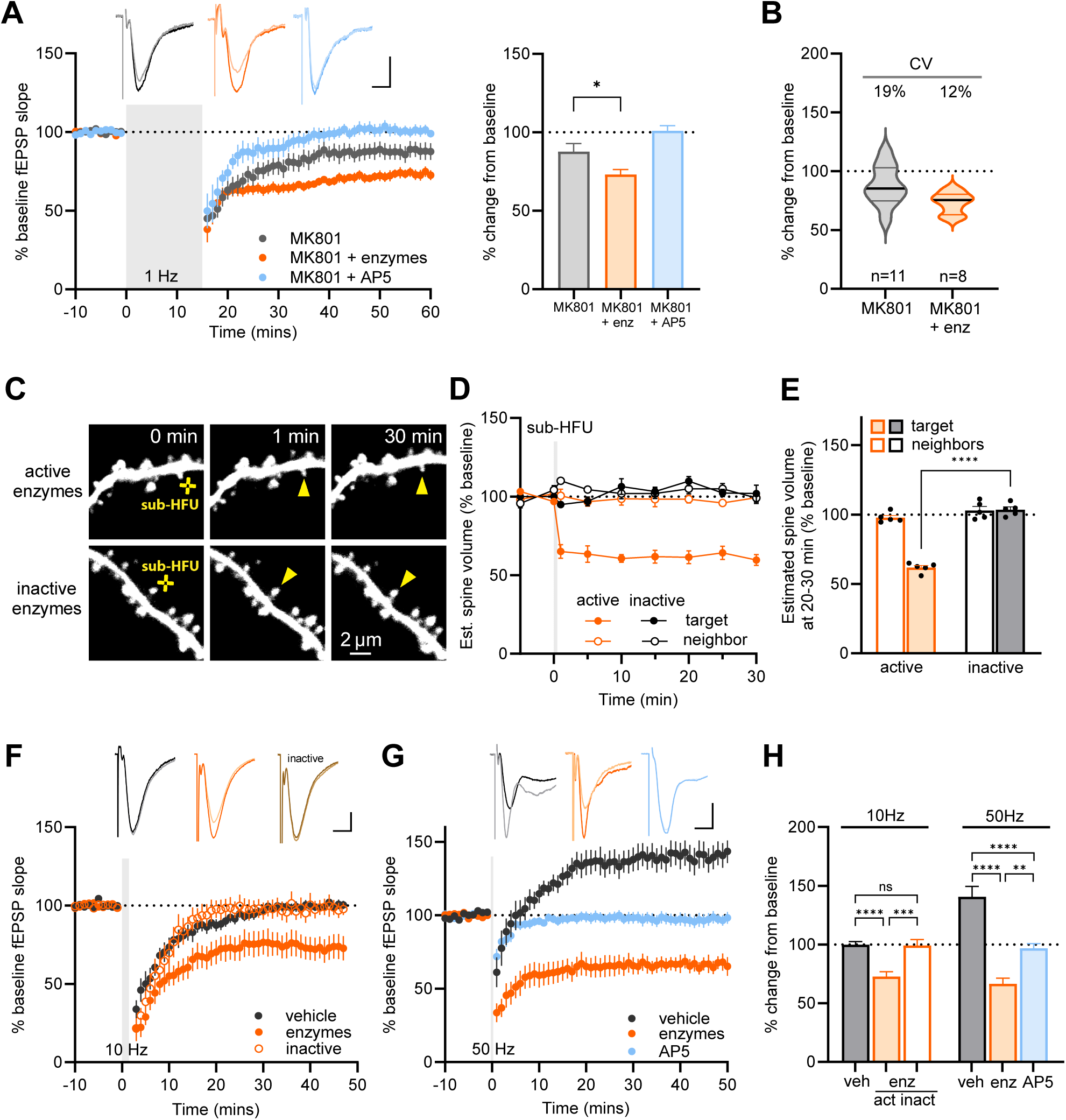
Enzymatic scavenging of endogenous co-agonists increases the magnitude and reduces the variance of ion flux-independent NMDAR-mediated LTD. **A**, *Left:* averaged plasticity experiments using a 1 Hz, 900 pulse LTD induction protocol in a recirculating bath with MK801 alone (grey circles; *n*=11), or in the presence of scavenging enzymes (orange circles; *n*=8) or 50 µM AP5 (light blue circles; *n*=6). *Right:* Compared with MK801 alone, the presence of scavenging enzymes resulted in an increased magnitude of non-ionotropic LTD (unpaired *t* test). **B,** Coefficient of variation (CV) of the experiments in A (*left*) and B (*right*) demonstrating the reduction in variation with either L689 or enzymes compared with MK801 alone. **C,** Images of dendrites of CA1 neurons from acute hippocampal slices GFP-M mice (P16-20) before and after sub-HFU (yellow crosses) at single spines across time (yellow arrowheads) in the presence of active scavenging enzymes (top) or heat-inactivated enzymes (bottom). **D-E,** Sub-HFU in the presence of active enzymes (orange filled circles/bar; 5 spines/5 cells), but not inactive enzymes (black filled circles/bar; 5 spines/5 cells), led to robust long-term spine shrinkage. Size of unstimulated neighbors (open circles/bars) did not change. **F,** averaged plasticity experiments using a neutral 10 Hz, 900 pulse protocol in a recirculating bath with vehicle (black circles; *n*=11), or in the presence of active scavenging enzymes (filled orange circles; *n*=9), or heat-inactivated enzymes (open orange circles; *n*=7). **G,** averaged plasticity experiments using a 50 Hz, 300 pulse LTP induction protocol in a recirculating bath with vehicle (black circles; *n*=10), or in the presence of scavenging enzymes (orange circles; *n*=10) or 50 µM AP5 (light blue circles; *n*=10). **H,** summary of data from C and D. One-way (electrophysiology data) and two-way (imaging data) ANOVA with Bonferroni post-hoc multiple comparisons test. Scale bars for all sample traces are 0.5 mV, 10 msec. All data represented as mean ± SEM. \*\**p*<0.01, \*\*\**p*<0.001, \*\*\*\**p*<0.0001, ^ns^*p*≥0.05.

In addition to LTD, ion flux-independent NMDAR signaling drives dendritic structural plasticity (Stein et al., 2015; Stein et al., 2020; Stein et al., 2021). We next tested whether scavenging of endogenous NMDAR co-agonists would also promote plasticity-induced long-term spine shrinkage. For these experiments, we chose to implement a neutral subthreshold high-frequency glutamate uncaging protocol (sub-HFU) that releases glutamate next to individual dendritic spines but does not result in any long-term spine size changes (Park et al., 2022b). We chose this protocol for its middle set point, which would allow detection of enzyme-induced changes in spine structural plasticity in either direction. Importantly, we found that following treatment with enzymes that scavenge endogenous co-agonists, but not with heat-inactivated enzymes, the sub-HFU protocol resulted in dendritic spine shrinkage (**Fig. 2C-E**; active enzymes: 62±2%, *n*=5; inactive enzymes, 103±2%, *n*=5; *p*<0.001, two-way ANOVA with Bonferroni post-hoc multiple comparisons test).

Similarly, a neutral 10 Hz electrical stimulation resulted in LTD when applied following enzymatic scavenging of endogenous co-agonists (**Fig. 2F,H** *left*; vehicle: 99.8±2.8%, *n*=11; enzymes: 72.7±4.2%, *n*=9; *p*<0.0001, one-way ANOVA with Bonferroni post-hoc multiple comparisons test) but not in the presence of heat-inactivated enzymes (**Fig. 2F,H** *left*; vehicle: 99.8±2.8%, *n*=11; inactivated enzymes: 99.1±5.1%, *n*=7; *p*=0.9992, one-way ANOVA with Bonferroni post-hoc multiple comparisons test).

Notably, a HFS of 50 Hz that typically induces long-term potentiation (LTP), also resulted in LTD following enzymatic scavenging of endogenous co-agonists (**Fig. 2G,H** *right*; vehicle: 140.7±8.8%, *n*=10; enzymes: 66.5±4.8%, *n*=10; AP5: 96.7±4.1%, *n*=10; *p*=0.0060 for enzymes vs. AP5, one-way ANOVA with Bonferroni post-hoc multiple comparisons test), providing evidence of the efficacy of the scavenging enzymes. Together, these results show that reducing co-agonist binding to the NMDAR, either via competitive antagonism or by decreasing co-agonist availability, promotes ion flux-independent NMDAR-mediated LTD and LTD-associated spine shrinkage.

### d-serine blocks ion flux-independent NMDAR-mediated LTD

To test the impact of increasing co-agonist site occupancy on ion flux-independent NMDAR-LTD, we next applied a saturating concentration (10 µM) of d-serine (Hansen et al., 2021). We again performed these experiments in the presence of MK801 and applied a standard 1 Hz, 900 stimuli low-frequency induction protocol. We found that 10 µM d-serine completely blocked LTD in the presence of MK801 (**Fig. 3A**; MK801: 80.2±4.0, *n*=12; +D-ser: 105.3±3.7, *n*=11; *p*=0.0002, unpaired *t* test), and reduced the inter-experiment variability (**Fig. 3B**). Importantly, d-serine application alone had no effect on fEPSP slope (**Fig 3C**; D-ser: 99.8±3.5% of baseline, *n*=7, *p*=0.9538, one sample *t* test).

**Figure 3.**
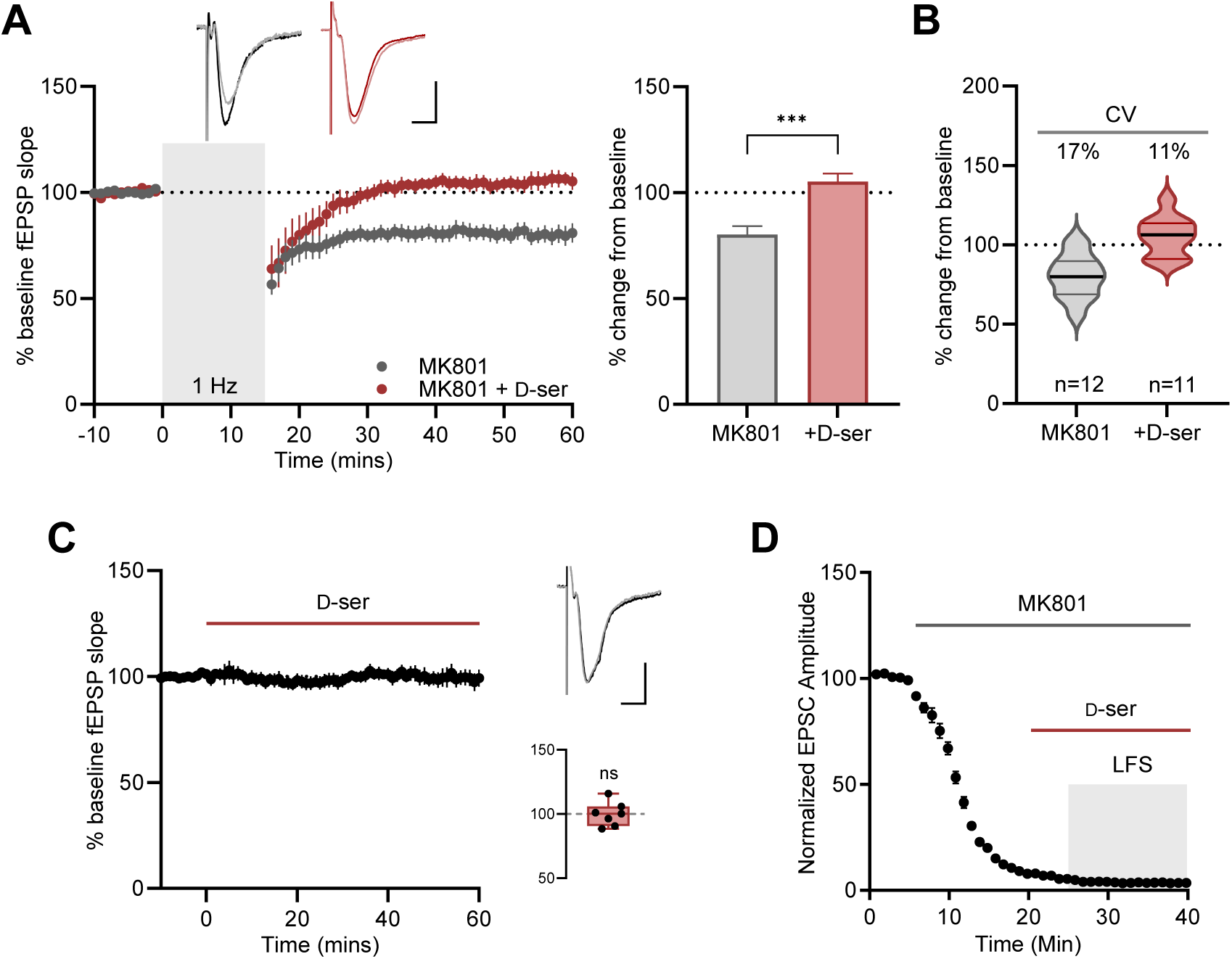
d-serine blocks ion flux-independent NMDAR-mediated LTD. **A,** *Left:* averaged plasticity experiments using a 1 Hz, 900 pulse LTD induction protocol in the presence of 100 µM MK801 alone (grey circles; *n*=12) or with the addition of 10 µM d-serine (red circles; *n*=11). *Right:* d-serine eliminated the non-ionotropic LTD observed with MK801 alone (unpaired *t* test). **B,** Coefficient of variation (CV) of the experiments in A demonstrating a reduction in variation with d-serine compared with MK801 alone. **C,** averaged fEPSP recordings normalized to the baseline fEPSP slope then 60 min incubation with 10 µM d-serine (*n*=7). *Inset:* d-serine did not change the fEPSP compared to baseline (one-sample *t* test). **D,** whole cell NMDAR-EPSCs experiment showing the time-course of MK801 block of synaptic receptors. 10 µM d-serine was added at 20 min and low frequency (1 Hz) stimulation (LFS) was started at 25 min. Neither d-serine incubation alone nor in combination with LFS altered the extent of MK801 blockade (*n*=10). Two-way ANOVA with Bonferroni post-hoc multiple comparisons test. Scale bars for all sample traces are 0.5 mV, 10 msec. All data represented as mean ± SEM. \*\**p*<0.01, \*\*\**p*<0.001.

MK801 has an extremely slow off-rate and has thus been considered to be effectively irreversible on the time scale of a typical experiment (Huettner and Bean, 1988; Reynolds and Miller, 1988; Jahr, 1992). However, the rate of MK801 dissociation from the NMDAR channel can be enhanced by agonist binding and channel re-opening (Huettner and Bean, 1988; MacDonald et al., 1991; McKay et al., 2013). Importantly, MK801 is continuously present in the bath during these experiments, and we found that during whole cell recordings of isolated NMDAR-EPSCs, neither 10 µM d-serine alone, nor in combination with low-frequency stimulation, resulted in any apparent transient or persistent changes in the degree of NMDAR inhibition by MK801 (**Fig 3D**).

### d-serine blocks LTD-associated spine shrinkage mediated by ion flux-independent NMDAR signaling

We next examined whether long-term dendritic spine shrinkage driven by ion flux-independent NMDAR signaling (Stein et al., 2015; Stein et al., 2020; Stein et al., 2021) was similarly blocked by d-serine. For these experiments, we used a low frequency uncaging protocol (LFU) designed to induce LTD and long-term spine shrinkage (Oh et al., 2013; Stein et al., 2020; Jang et al., 2021). We isolated ion flux-independent NMDAR-mediated plasticity by examining spine shrinkage with an LTD-inducing stimulus in the presence of MK801. Indeed, we found that addition of 10 µM d-serine completely blocked long-term spine shrinkage induced by LFU at single dendritic spines in the presence of MK801 (**Fig. 4A-C**; veh: 69±2%, *n*=6; +D-Ser, 107±9%, *n*=7; *p*<0.001, two-way ANOVA with Bonferroni post-hoc multiple comparisons test). No changes in size of neighboring spines were observed. Thus, addition of d-serine is sufficient to completely inhibit LTD-associated long-term dendritic spine shrinkage mediated by ion flux-independent NMDAR signaling.

**Figure 4.**
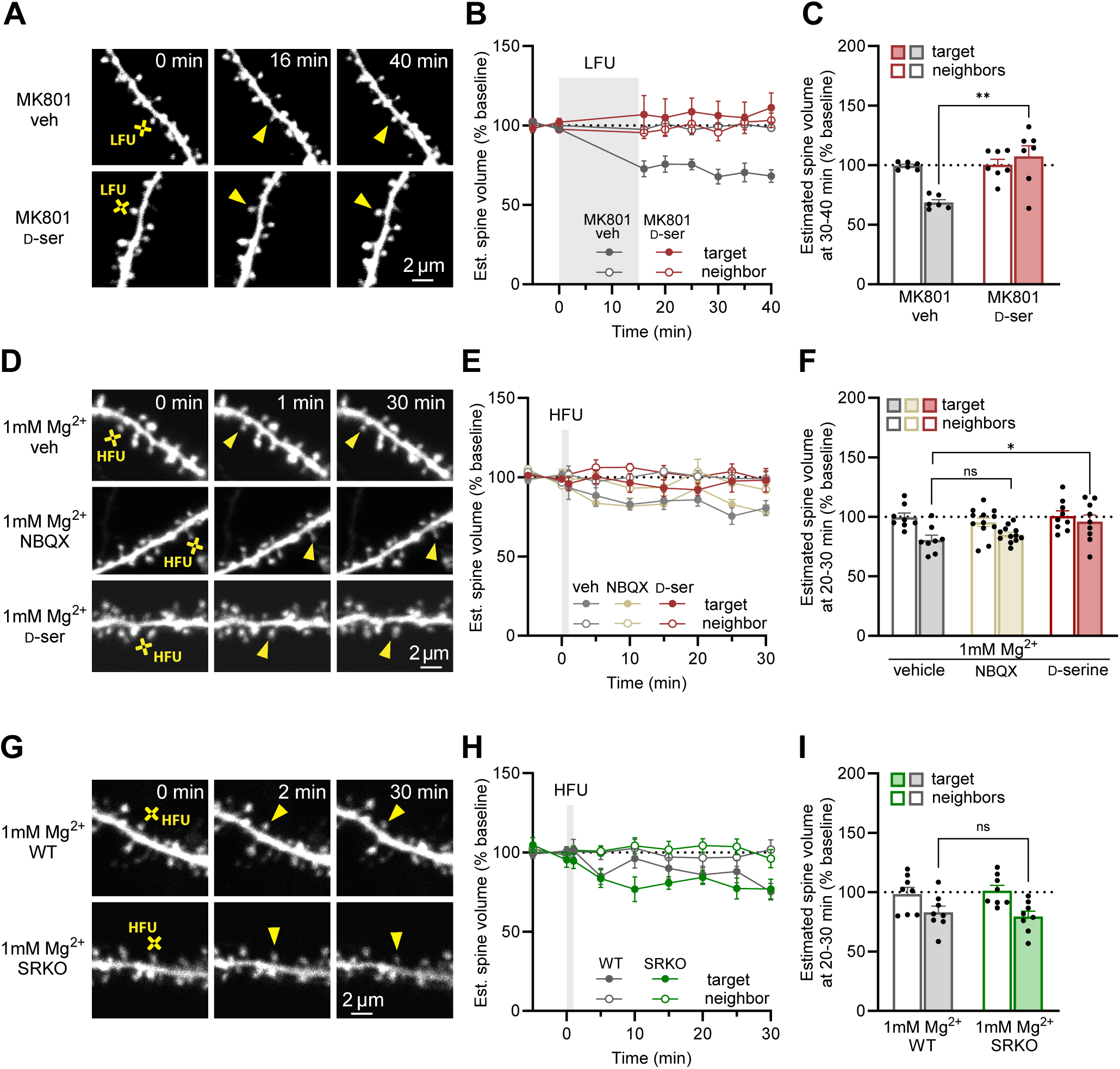
d-serine blocks LTD-associated spine shrinkage mediated by ion flux-independent NMDAR signaling. **A,** Images of dendrites of CA1 neurons from acute hippocampal slices GFP-M mice (P16-20) before and after LFU (yellow crosses) at single spines across time (yellow arrowheads) in the presence of MK-801 and vehicle (top) or d-serine (10 µM; bottom). **B-C,** LFU in the presence of 100 µM MK801 (black filled circles/bar; 6 spines/6 cells) led to robust spine shrinkage, which was fully inhibited with the addition of d-serine (red filled circles/bar; 7 spines/7 cells). Size of unstimulated neighbors (open circles/bars) did not change. **D,** Images of dendrites of CA1 neurons in acute hippocampal slices from GFP-M mice (P17-21) before and after HFU (yellow crosses) at single spines in 1 mM Mg^2+^ and either vehicle (water; top) NBQX (50 µM; middle) or d-serine (10 µM; bottom). **E-F**, HFU-induced long-term spine shrinkage in 1 mM Mg^2+^ was blocked by d-serine (red filled circles/bar; 9 spines/9 cells), but not by vehicle (black filled circles/bar; 8 spines/8 cells) or by NBQX (beige filled circles/bar; 12 spines/12 cells). Size of unstimulated neighbors (open circles/bars) did not change. **G,** Images of dendrites of CA1 neurons in acute hippocampal slices from WT/GFP-M (top) and SRKO/GFP-M (bottom) littermates (P17-21) before and after HFU (yellow crosses) at single spines in 1 mM Mg^2+^. **H-I,** Both WT (black filled circles/bar; 8 spines/8 cells) and SRKO (purple filled circles/bar; 8 spines/8 cells) exhibited robust HFU-induced long-term spine shrinkage in 1 mM Mg^2+^. Size of unstimulated neighbors (open circles/bars) did not change. Two-way ANOVA with Bonferroni post-hoc multiple comparisons test. All data represented as mean ± SEM. \**p*<0.01, \*\**p*<0.001, ^ns^*p*>0.05.

Physiologically, ion flux-independent NMDAR-mediated forms of plasticity and signaling are likely to be most relevant at individual synapses during Mg^2+^ block of ion flow through the NMDAR, as expected at resting membrane potentials and at silent synapses, or in physiological or disease states associated with reduced d-serine levels (Park et al., 2022b; Park et al., 2022a). Because our single-spine plasticity protocols were originally developed using low or zero Mg^2+^ to mimic coincident pre- and post-synaptic activity (Oh et al., 2013; Jang et al., 2021), we chose to examine next whether d-serine levels would act to modulate NMDAR-mediated structural plasticity during Mg^2+^ block. For these experiments, we implemented a high frequency glutamate uncaging protocol (HFU) that typically results in long-term potentiation (LTP) and spine growth but, in the presence of MK801 or L689 to block ion flow through the NMDAR, instead leads to a robust LTD and LTD-associated spine shrinkage mediated by ion flux-independent NMDAR signaling (Stein et al., 2015; Stein et al., 2020; Stein et al., 2021).

In the presence of 1 mM Mg^2+^, we observed that HFU at individual dendritic spines resulted in long-term spine shrinkage, as expected from ion flux-independent NMDAR signaling (**Fig. 4D-F**; veh: 81±4%, *n*=8, *p*<0.001 compared to baseline, two-way ANOVA with Bonferroni post-hoc multiple comparisons test). Importantly, the addition of 10 µM d-serine completely blocked HFU-induced long-term spine shrinkage in the presence of 1 mM Mg^2+^ (**Fig. 4D-F**; D-Ser: 96±4%, *n*=9, *p*=0.001 compared to vehicle, two-way ANOVA with Bonferroni post-hoc multiple comparisons test). Notably, the magnitude of spine shrinkage observed in the presence of 1 mM Mg^2+^ was comparable to that observed with the addition of 50 µM NBQX to inhibit AMPARs (**Fig. 4D-F**; NBQX: 85±3%, *n*=12, *p*=0.75, two-way ANOVA with Bonferroni post-hoc multiple comparisons test) and in the SRKO mouse that lacks the enzyme for d-serine production (**Fig. 4G-I**; WT littermates: 83±5%, *n*=8; SRKO: 80±5%, *n*=8; *p*>0.99, two-way ANOVA with Bonferroni post-hoc multiple comparisons test), supporting that spine shrinkage observed in the presence of 1 mM Mg^2+^ is mediated by ion flux-independent NMDAR signaling and thus that the block of spine shrinkage is not due to d-serine-mediated enhancement of ion flux through the NMDAR. Therefore, we conclude that addition of d-serine is sufficient to block long-term dendritic spine shrinkage mediated by ion flux-independent NMDAR signaling under physiological conditions expected at resting membrane potentials and at silent synapses.

### d-serine inhibits NMDA-induced intracellular GluN1 conformational changes

Previous work using FRET-FLIM (Förster resonance energy transfer - fluorescence lifetime imaging microscopy) has demonstrated that glutamate or NMDA binding can induce conformational movements in the NMDAR intracellular c-terminal domain in the presence of co-agonist site antagonists or MK801 (Dore et al., 2015). Thus, NMDARs can transmit agonist-driven information into the cell in the absence of ion flux through the receptor channel. Using this FRET-FLIM approach, we found that 10 µM d-serine completely inhibits the NMDA-induced lifetime change in dendritic spines (**Fig. 5**; MK801: 88±26ps, *n*=20 neurons; +D-ser: -7±13ps, *n*=23 neurons; *p*=0.0015, unpaired *t* test). Together, these results suggest that d-serine inhibits ion flux-independent NMDAR-mediated LTD and long-term spine shrinkage by preventing intracellular conformational changes of the NMDAR in response to glutamate.

**Figure 5.**
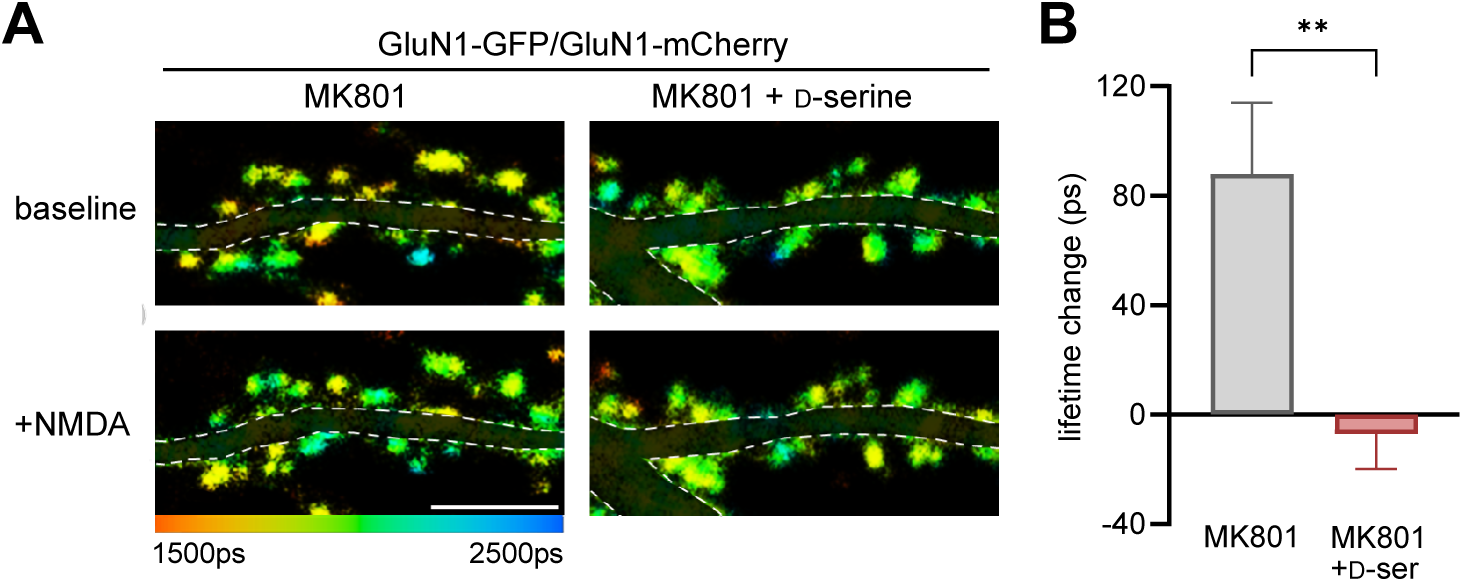
d-serine inhibits NMDA-induced intracellular GluN1 conformational changes. **A,** Representative fluorescence lifetime images of neurons expressing GluN1-GFP and GluN1-mCherry (with GluN2B) before and during treatment with 25 µM NMDA. Neurons were incubated with MK801 alone (left) or with 10 µM d-serine (right). Pseudocolor scale indicates GFP lifetime at each pixel. Scale bar, 5 µm; dendritic segments masked for clarity. **B,** Average NMDA-induced spine GluN1-GFP lifetime change for indicated conditions, *N* (>20 neurons, >500 spines) for each condition, showing that 10 µM d-serine completely blocks the NMDA-induced lifetime change, unpaired *t* test. Data represented as mean ± SEM. \*\**p*<0.01.

## Discussion

Over the past decade, evidence has accumulated supporting that ion flux-independent (non-ionotropic) signaling of the NMDAR is sufficient to drive synaptic plasticity. Indeed, studies from several independent laboratories have shown that glutamate binding to NMDARs, independent of ion flux, is sufficient to induce LTD (Nabavi et al., 2013; Stein et al., 2015; Carter and Jahr, 2016; Wong and Gray, 2018; Stein et al., 2020; Dore and Malinow, 2021) and LTD-associated dendritic spine shrinkage (Birnbaum et al., 2015; Stein et al., 2015; Stein et al., 2020; Stein et al., 2021; Thomazeau et al., 2021; Park et al., 2022b). Physiologically, such ion flux-independent signaling through NMDARs would be expected with glutamate binding (1) when levels of co-agonist are low, such as could occur during sleep (e.g. Papouin et al., 2017) and in diseases associated with lowered d-serine levels (e.g. Park et al., 2022b), or (2) when the NMDAR channel pore is blocked by Mg^2+^, such as at silent synapses lacking AMPARs or when there is insufficient postsynaptic depolarization to relieve the Mg^2+^ block.

Here, we report that even in the latter case when the NMDAR pore is blocked, occupancy of the NMDAR co-agonist binding site is a major determinant of ion flux-independent plasticity. Specifically, using MK801 or Mg^2+^ to isolate ion flux-independent signaling, we found that increasing d-serine to levels that would be expected to saturate the NMDAR co-agonist binding site completely inhibited ion flux-independent NMDAR-mediated LTD and LTD associated spine shrinkage. Thus, altered levels of d-serine, either through physiological or pathological conditions, will not only determine, but will regulate the extent of ion flux-independent NMDAR signaling.

How might occupancy of the NMDAR co-agonist site interfere with plasticity mediated by ion flux-independent NMDAR signaling? Here we show that d-serine inhibits NMDA-induced conformational changes of the intracellular C-tails of the NMDAR as measured by FRET-FLIM. Several studies have examined the conformational movements of the NMDAR during non-ionotropic signaling using FRET-FLIM to monitor movements of intracellular carboxy-terminal tails (C-tails) of the GluN1 subunits (Aow et al., 2015; Dore et al., 2015). In the original reports (Aow et al., 2015; Dore et al., 2015), glutamate (or NMDA) binding in the presence of MK801 or L689 resulted in an increased GFP lifetime, demonstrating that the GluN1 C-tails move away from each other during ion flux-independent signaling. In the current study, this NMDA-induced movement is completely blocked by d-serine. Interestingly, in a study by another group, d-serine binding alone (i.e. in the absence of NMDA or glutamate) shortened the GFP lifetime suggesting that it causes the GluN1 C-tails to move closer together (Ferreira et al., 2017), opposite to the effect of glutamate. Structurally, a dimer-dimer interface between the ligand binding domains of GluN1 and GluN2 (Furukawa et al., 2005) results in allosteric coupling between glutamate and co-agonist binding (Regalado et al., 2001; Cummings and Popescu, 2015; Durham et al., 2020) providing a potential structural mechanism for the inhibition of non-ionotropic NMDAR signaling by d-serine.

NMDARs are well-suited to employ conformation-based signaling. Compared with AMPA receptors, NMDARs adopt a significantly more compact structure of their extracellular domains (Karakas and Furukawa, 2014; Lee et al., 2014), which explains their wide-range of allosteric modulation as this compact structure provides molecular routes for the transmission of conformational changes through the complex. However, the large intracellular C-terminal domains of NMDARs are considered intrinsically unstructured which would seemingly limit transmission of conformational signals intracellularly. In addition to the FRET-FLIM studies, multiple lines of evidence support conformational interactions between the C-terminal and the other domains of NMDARs. For example, deletion or truncation of GluN2 C-terminal domains has been shown increase co-agonist potency (Puddifoot et al., 2009), reduce single-channel open probability (Maki et al., 2012; Punnakkal et al., 2012), and affect the activity of extracellular allosteric modulators of NMDARs (Sapkota et al., 2019). Phosphorylation of individual residues on GluN2 C-terminal tails by either PKC or PKA increases NMDAR single-channel open probability (Zheng et al., 1999; Lan et al., 2001; Aman et al., 2014; Murphy et al., 2014) and alters allosteric modulator activity (Sapkota et al., 2019), suggesting transmission of intracellular changes to the core channel domains of the receptor. Furthermore, proteins that interact with the intracellular C-terminal domains of NMDARs have been shown to affect channel gating. Co-expression of PSD-95 along with NMDARs in heterologous cells enhances single-channel opening rate and reduces agonist potency (Yamada et al., 1999; Lin et al., 2004) and Ca^2+^-calmodulin binding to the GluN1 C-terminal domain affects channel gating properties (Zhang et al., 1998; Iacobucci and Popescu, 2017). Thus, while the C-terminal domains of NMDARs are intrinsically disordered, synaptic NMDARs are a part of a large multi-protein complex at the postsynaptic density (Fan et al., 2014; Frank et al., 2016; Frank and Grant, 2017) that likely imparts the secondary and tertiary structure required to transmit information via agonist-induced conformational changes.

Despite the growing evidence supporting ion flux-independent signaling through the NMDAR, significant controversy has remained. Most of the contradictory results regarding ion flux-independent LTD have been with the use of MK801 to inhibit ion flux through the pore. In the presence of MK801, some groups observed robust LTD and LTD-associated spine shrinkage (Mayford et al., 1995; Nabavi et al., 2013; Dore et al., 2015; Stein et al., 2015; Carter and Jahr, 2016), while others saw no LTD (Sanderson et al., 2012; Babiec et al., 2014; Coultrap et al., 2014; Sanderson et al., 2016). Importantly, while MK801 effectively blocks ion-flux through the NMDAR channel, it does not significantly alter the affinity or occupancy of the co-agonist binding site (MacDonald et al., 1991). Uncompetitive NMDAR open-channel blockers like MK801 are considered “trapping” blockers, entering the pore of the channel when it is open then promoting the return of the receptor to the closed state, while returning to inactive conformations that allow for the release of glutamate and d-serine (Huettner and Bean, 1988; Jahr and Stevens, 1990; MacDonald et al., 1991; Blanpied et al., 1997; Bolshakov et al., 2003; Song et al., 2018). Here we report that LTD in the presence of the open channel blocker MK801 shows a higher inter-experiment variability than observed when the occupancy of the NMDAR co-agonist site is directly controlled by saturating it with d-serine, blocking it with L689, or greatly reducing it with co-agonist scavenging enzymes. We suggest that experimental differences in d-serine availability could serve as a source of the inconsistencies observed when using MK801. Various factors could affect the availability of d-serine in slice preparations, including but not limited to, age, strain, slice health, preparation method, slice orientation, slice thickness, and perfusion rate. Alternatively, MK801, as an open channel blocker, might not be consistently effective in its block of ion flux due to the requirement for receptor activation and channel opening prior to blockade (Huettner and Bean, 1988); however, we and other have shown this is unlikely (Nabavi et al., 2013; Stein et al., 2015; Carter and Jahr, 2016).

Physiologically, ion flux-independent NMDAR-mediated plasticity is likely to play a key role during development, where it could serve to modulate the stability of immature ‘silent’ synapses that lack postsynaptic AMPA receptors. At silent synapses, glutamate release alone would be insufficient to invoke the local depolarization needed to remove the NMDAR Mg^2+^ block. Therefore, glutamate binding to NMDARs at silent synapses would be predicted to primarily engage ion flux-independent NMDAR-mediated LTD, representing a mechanism to continuously exclude AMPA receptors and maintain synapse silence (Colonnese et al., 2003; Xiao et al., 2004; Colonnese et al., 2005; Ultanir et al., 2007; Adesnik et al., 2008; Kerchner and Nicoll, 2008; Gray et al., 2011), and to drive shrinkage and loss of synapses that do not exhibit coincident pre- and postsynaptic activity (Feldman, 2012). Indeed, previous studies have also established that holding neurons at hyperpolarized potentials during low-frequency stimulation does not inhibit LTD (Nabavi et al., 2013) and leads to a modest synaptic depression following a normally LTP-inducing high-frequency stimulation (Malinow and Miller, 1986).

Notably, we found that NMDAR-mediated spine shrinkage in the presence of Mg^2+^, a condition expected physiologically during a weak stimulus without co-incident postsynaptic activity, is inhibited by d-serine. Intriguingly, serine racemase expression in CA1 pyramidal neurons increases between the second and fourth postnatal weeks (Miya et al., 2008; Folorunso et al., 2021), which corresponds with the increased usage of d-serine as a synaptic NMDAR co-agonist (Le Bail et al., 2015). Thus, increased d-serine could serve to reduce the prevalence of ion flux-independent NMDAR-mediated plasticity following the critical period where excessive synapse pruning matches the developing nervous system to the sensory environment.

## Acknowledgements

This work was supported by the NIH (R01MH117130 and R21MH116315 to JAG, R01NS062736 to KZ, RF1AG0677049 to KD, and T32GM099608 to EVB, and F31NS122488 to SP) and the Deutsche Forschungsgemeinschaft Walter Benjamin project 468470832 (to MA). Thank you to Herman Wolosker (Technion – Israel Institute of Technology) for generously gifting the enzymes.

## Author contributions

JAG conceived the initial study; EVB, MA, KD, KZ, and JG designed research; EVB, MA, VV, ACC, DKP, RJS, AFN, and SP performed research and analyzed data; JAG and KZ wrote the paper; all authors edited the paper.

## References

Adesnik H, Li G, During MJ, Pleasure SJ, Nicoll RA (2008) NMDA receptors inhibit synapse unsilencing during brain development. Proc Natl Acad Sci U S A 105:5597–5602.

Aman TK, Maki BA, Ruffino TJ, Kasperek EM, Popescu GK (2014) Separate intramolecular targets for protein kinase A control N-methyl-D-aspartate receptor gating and Ca2+ permeability. J Biol Chem 289:18805–18817.

Aow J, Dore K, Malinow R (2015) Conformational signaling required for synaptic plasticity by the NMDA receptor complex. Proc Natl Acad Sci U S A 112:14711–14716.

Babiec WE, Guglietta R, Jami SA, Morishita W, Malenka RC, O’Dell TJ (2014) Ionotropic NMDA receptor signaling is required for the induction of long-term depression in the mouse hippocampal CA1 region. J Neurosci 34:5285–5290.

Basu AC, Tsai GE, Ma CL, Ehmsen JT, Mustafa AK, Han L, Jiang ZI, Benneyworth MA, Froimowitz MP, Lange N, Snyder SH, Bergeron R, Coyle JT (2009) Targeted disruption of serine racemase affects glutamatergic neurotransmission and behavior. Mol Psychiatry 14:719–727.

Birnbaum JH, Bali J, Rajendran L, Nitsch RM, Tackenberg C (2015) Calcium flux-independent NMDA receptor activity is required for Abeta oligomer-induced synaptic loss. Cell Death Dis 6:e1791.

Blanpied TA, Boeckman FA, Aizenman E, Johnson JW (1997) Trapping channel block of NMDA-activated responses by amantadine and memantine. J Neurophysiol 77:309–323.

Bolshakov KV, Gmiro VE, Tikhonov DB, Magazanik LG (2003) Determinants of trapping block of N-methyl-d-aspartate receptor channels. J Neurochem 87:56–65.

Carter BC, Jahr CE (2016) Postsynaptic, not presynaptic NMDA receptors are required for spike-timing-dependent LTD induction. Nat Neurosci 19:1218–1224.

Colonnese MT, Shi J, Constantine-Paton M (2003) Chronic NMDA receptor blockade from birth delays the maturation of NMDA currents, but does not affect AMPA/kainate currents. J Neurophysiol 89:57–68.

Colonnese MT, Zhao JP, Constantine-Paton M (2005) NMDA receptor currents suppress synapse formation on sprouting axons in vivo. J Neurosci 25:1291–1303.

Coultrap SJ, Freund RK, O’Leary H, Sanderson JL, Roche KW, Dell’Acqua ML, Bayer KU (2014) Autonomous CaMKII mediates both LTP and LTD using a mechanism for differential substrate site selection. Cell Rep 6:431–437.

Cummings KA, Popescu GK (2015) Glycine-dependent activation of NMDA receptors. J Gen Physiol 145:513–527.

Dore K, Malinow R (2021) Elevated PSD-95 Blocks Ion-flux Independent LTD: A Potential New Role for PSD-95 in Synaptic Plasticity. Neuroscience 456:43–49.

Dore K, Aow J, Malinow R (2015) Agonist binding to the NMDA receptor drives movement of its cytoplasmic domain without ion flow. Proc Natl Acad Sci U S A 112:14705–14710.

Dore K, Aow J, Malinow R (2016) The Emergence of NMDA Receptor Metabotropic Function: Insights from Imaging. Front Synaptic Neurosci 8:20.

Dudek SM, Bear MF (1992) Homosynaptic long-term depression in area CA1 of hippocampus and effects of N-methyl-D-aspartate receptor blockade. Proc Natl Acad Sci U S A 89:4363–4367.

Dupourque D, Newton WA, Snell EE (1966) Purification and properties of d-serine dehydrase from Escherichia coli. J Biol Chem 241:1233–1238.

Durham RJ, Paudyal N, Carrillo E, Bhatia NK, Maclean DM, Berka V, Dolino DM, Gorfe AA, Jayaraman V (2020) Conformational spread and dynamics in allostery of NMDA receptors. Proc Natl Acad Sci U S A 117:3839–3847.

Fan X, Jin WY, Wang YT (2014) The NMDA receptor complex: a multifunctional machine at the glutamatergic synapse. Front Cell Neurosci 8:160.

Feldman DE (2012) The spike-timing dependence of plasticity. Neuron 75:556–571.

Feng G, Mellor RH, Bernstein M, Keller-Peck C, Nguyen QT, Wallace M, Nerbonne JM, Lichtman JW, Sanes JR (2000) Imaging neuronal subsets in transgenic mice expressing multiple spectral variants of GFP. Neuron 28:41–51.

Ferreira JS, Papouin T, Ladepeche L, Yao A, Langlais VC, Bouchet D, Dulong J, Mothet JP, Sacchi S, Pollegioni L, Paoletti P, Oliet SHR, Groc L (2017) Co-agonists differentially tune GluN2B-NMDA receptor trafficking at hippocampal synapses. Elife 6.

Folorunso OO, Harvey TL, Brown SE, Cruz C, Shahbo E, Ajjawi I, Balu DT (2021) Forebrain expression of serine racemase during postnatal development. Neurochem Int 145:104990.

Frank RA, Grant SG (2017) Supramolecular organization of NMDA receptors and the postsynaptic density. Curr Opin Neurobiol 45:139–147.

Frank RA, Komiyama NH, Ryan TJ, Zhu F, O’Dell TJ, Grant SG (2016) NMDA receptors are selectively partitioned into complexes and supercomplexes during synapse maturation. Nat Commun 7:11264.

Furukawa H, Singh SK, Mancusso R, Gouaux E (2005) Subunit arrangement and function in NMDA receptors. Nature 438:185–192.

Gray JA, Zito K, Hell JW (2016) Non-ionotropic signaling by the NMDA receptor: controversy and opportunity. F1000Res 5.

Gray JA, Shi Y, Usui H, During MJ, Sakimura K, Nicoll RA (2011) Distinct modes of AMPA receptor suppression at developing synapses by GluN2A and GluN2B: single-cell NMDA receptor subunit deletion in vivo. Neuron 71:1085–1101.

Gustafson EC, Stevens ER, Wolosker H, Miller RF (2007) Endogenous d-serine contributes to NMDA-receptor-mediated light-evoked responses in the vertebrate retina. J Neurophysiol 98:122–130.

Hansen KB, Yi F, Perszyk RE, Furukawa H, Wollmuth LP, Gibb AJ, Traynelis SF (2018) Structure, function, and allosteric modulation of NMDA receptors. J Gen Physiol 150:1081–1105.

Hansen KB, Wollmuth LP, Bowie D, Furukawa H, Menniti FS, Sobolevsky AI, Swanson GT, Swanger SA, Greger IH, Nakagawa T, McBain CJ, Jayaraman V, Low CM, Dell’Acqua ML, Diamond JS, Camp CR, Perszyk RE, Yuan H, Traynelis SF (2021) Structure, Function, and Pharmacology of Glutamate Receptor Ion Channels. Pharmacol Rev 73:298-487.

Huettner JE, Bean BP (1988) Block of N-methyl-D-aspartate-activated current by the anticonvulsant MK-801: selective binding to open channels. Proc Natl Acad Sci U S A 85:1307–1311.

Iacobucci GJ, Popescu GK (2017) Resident Calmodulin Primes NMDA Receptors for Ca(2+)-Dependent Inactivation. Biophys J 113:2236–2248.

Jahr CE (1992) High probability opening of NMDA receptor channels by L-glutamate. Science 255:470–472.

Jahr CE, Stevens CF (1990) A quantitative description of NMDA receptor-channel kinetic behavior. J Neurosci 10:1830–1837.

Jang J, Anisimova M, Oh WC, Zito K (2021) Induction of input-specific spine shrinkage on dendrites of rodent hippocampal CA1 neurons using two-photon glutamate uncaging. STAR Protoc 2:100996.

Job V, Molla G, Pilone MS, Pollegioni L (2002) Overexpression of a recombinant wild-type and His-tagged Bacillus subtilis glycine oxidase in Escherichia coli. Eur J Biochem 269:1456–1463.

Karakas E, Furukawa H (2014) Crystal structure of a heterotetrameric NMDA receptor ion channel. Science 344:992–997.

Kartvelishvily E, Shleper M, Balan L, Dumin E, Wolosker H (2006) Neuron-derived d-serine release provides a novel means to activate N-methyl-D-aspartate receptors. J Biol Chem 281:14151–14162.

Kerchner GA, Nicoll RA (2008) Silent synapses and the emergence of a postsynaptic mechanism for LTP. Nat Rev Neurosci 9:813–825.

Lan JY, Skeberdis VA, Jover T, Grooms SY, Lin Y, Araneda RC, Zheng X, Bennett MV, Zukin RS (2001) Protein kinase C modulates NMDA receptor trafficking and gating. Nat Neurosci 4:382–390.

Le Bail M, Martineau M, Sacchi S, Yatsenko N, Radzishevsky I, Conrod S, Ait Ouares K, Wolosker H, Pollegioni L, Billard JM, Mothet JP (2015) Identity of the NMDA receptor coagonist is synapse specific and developmentally regulated in the hippocampus. Proc Natl Acad Sci U S A 112:E204–213.

Lee CH, Lu W, Michel JC, Goehring A, Du J, Song X, Gouaux E (2014) NMDA receptor structures reveal subunit arrangement and pore architecture. Nature 511:191–197.

Li Y, Sacchi S, Pollegioni L, Basu AC, Coyle JT, Bolshakov VY (2013) Identity of endogenous NMDAR glycine site agonist in amygdala is determined by synaptic activity level. Nat Commun 4:1760.

Lin Y, Skeberdis VA, Francesconi A, Bennett MV, Zukin RS (2004) Postsynaptic density protein-95 regulates NMDA channel gating and surface expression. J Neurosci 24:10138–10148.

Lu JM, Gong N, Wang YC, Wang YX (2012) D-Amino acid oxidase-mediated increase in spinal hydrogen peroxide is mainly responsible for formalin-induced tonic pain. Br J Pharmacol 165:1941–1955.

MacDonald JF, Bartlett MC, Mody I, Pahapill P, Reynolds JN, Salter MW, Schneiderman JH, Pennefather PS (1991) Actions of ketamine, phencyclidine and MK-801 on NMDA receptor currents in cultured mouse hippocampal neurones. J Physiol 432:483–508.

Maki BA, Aman TK, Amico-Ruvio SA, Kussius CL, Popescu GK (2012) C-terminal domains of N-methyl-D-aspartic acid receptor modulate unitary channel conductance and gating. J Biol Chem 287:36071–36080.

Malinow R, Miller JP (1986) Postsynaptic hyperpolarization during conditioning reversibly blocks induction of long-term potentiation. Nature 320:529–530.

Matlashov ME, Belousov VV, Enikolopov G (2014) How much H(2)O(2) is produced by recombinant D-amino acid oxidase in mammalian cells? Antioxid Redox Signal 20:1039–1044.

Mayford M, Wang J, Kandel ER, O’Dell TJ (1995) CaMKII regulates the frequency-response function of hippocampal synapses for the production of both LTD and LTP. Cell 81:891–904.

McKay S, Bengtson CP, Bading H, Wyllie DJ, Hardingham GE (2013) Recovery of NMDA receptor currents from MK-801 blockade is accelerated by Mg2+ and memantine under conditions of agonist exposure. Neuropharmacology 74:119–125.

Meunier CN, Dallerac G, Le Roux N, Sacchi S, Levasseur G, Amar M, Pollegioni L, Mothet JP, Fossier P (2016) d-serine and Glycine Differentially Control Neurotransmission during Visual Cortex Critical Period. PLoS One 11:e0151233.

Miya K, Inoue R, Takata Y, Abe M, Natsume R, Sakimura K, Hongou K, Miyawaki T, Mori H (2008) Serine racemase is predominantly localized in neurons in mouse brain. J Comp Neurol 510:641–654.

Mothet JP, Parent AT, Wolosker H, Brady RO, Jr., Linden DJ, Ferris CD, Rogawski MA, Snyder SH (2000) d-serine is an endogenous ligand for the glycine site of the N-methyl-D-aspartate receptor. Proc Natl Acad Sci U S A 97:4926–4931.

Murphy JA, Stein IS, Lau CG, Peixoto RT, Aman TK, Kaneko N, Aromolaran K, Saulnier JL, Popescu GK, Sabatini BL, Hell JW, Zukin RS (2014) Phosphorylation of Ser1166 on GluN2B by PKA is critical to synaptic NMDA receptor function and Ca2+ signaling in spines. J Neurosci 34:869–879.

Nabavi S, Kessels HW, Alfonso S, Aow J, Fox R, Malinow R (2013) Metabotropic NMDA receptor function is required for NMDA receptor-dependent long-term depression. Proc Natl Acad Sci U S A 110:4027–4032.

Oh WC, Hill TC, Zito K (2013) Synapse-specific and size-dependent mechanisms of spine structural plasticity accompanying synaptic weakening. Proc Natl Acad Sci U S A 110:E305–312.

Panatier A, Theodosis DT, Mothet JP, Touquet B, Pollegioni L, Poulain DA, Oliet SH (2006) Glia-derived d-serine controls NMDA receptor activity and synaptic memory. Cell 125:775–784.

Papouin T, Dunphy JM, Tolman M, Dineley KT, Haydon PG (2017) Septal Cholinergic Neuromodulation Tunes the Astrocyte-Dependent Gating of Hippocampal NMDA Receptors to Wakefulness. Neuron 94:840–854 e847.

Papouin T, Ladepeche L, Ruel J, Sacchi S, Labasque M, Hanini M, Groc L, Pollegioni L, Mothet JP, Oliet SH (2012) Synaptic and extrasynaptic NMDA receptors are gated by different endogenous coagonists. Cell 150:633–646.

Park DK, Stein IS, Zito K (2022a) Ion flux-independent NMDA receptor signaling. Neuropharmacology 210:109019.

Park DK, Petshow S, Anisimova M, Barragan EV, Gray JA, Stein IS, Zito K (2022b) Reduced d-serine levels drive enhanced non-ionotropic NMDA receptor signaling and destabilization of dendritic spines in a mouse model for studying schizophrenia. Neurobiol Dis 170:105772.

Puddifoot CA, Chen PE, Schoepfer R, Wyllie DJ (2009) Pharmacological characterization of recombinant NR1/NR2A NMDA receptors with truncated and deleted carboxy termini expressed in Xenopus laevis oocytes. Br J Pharmacol 156:509–518.

Punnakkal P, Jendritza P, Kohr G (2012) Influence of the intracellular GluN2 C-terminal domain on NMDA receptor function. Neuropharmacology 62:1985–1992.

Regalado MP, Villarroel A, Lerma J (2001) Intersubunit cooperativity in the NMDA receptor. Neuron 32:1085–1096.

Reynolds IJ, Miller RJ (1988) Multiple sites for the regulation of the N-methyl-D-aspartate receptor. Mol Pharmacol 33:581–584.

Rosini E, Piubelli L, Molla G, Frattini L, Valentino M, Varriale A, D’Auria S, Pollegioni L (2014) Novel biosensors based on optimized glycine oxidase. FEBS J 281:3460–3472.

Sanderson JL, Gorski JA, Dell’Acqua ML (2016) NMDA Receptor-Dependent LTD Requires Transient Synaptic Incorporation of Ca(2)(+)-Permeable AMPARs Mediated by AKAP150-Anchored PKA and Calcineurin. Neuron 89:1000–1015.

Sanderson JL, Gorski JA, Gibson ES, Lam P, Freund RK, Chick WS, Dell’Acqua ML (2012) AKAP150-anchored calcineurin regulates synaptic plasticity by limiting synaptic incorporation of Ca2+-permeable AMPA receptors. J Neurosci 32:15036–15052.

Sapkota K, Dore K, Tang K, Irvine M, Fang G, Burnell ES, Malinow R, Jane DE, Monaghan DT (2019) The NMDA receptor intracellular C-terminal domains reciprocally interact with allosteric modulators. Biochem Pharmacol 159:140–153.

Settembre EC, Dorrestein PC, Park JH, Augustine AM, Begley TP, Ealick SE (2003) Structural and mechanistic studies on ThiO, a glycine oxidase essential for thiamin biosynthesis in Bacillus subtilis. Biochemistry 42:2971–2981.

Shleper M, Kartvelishvily E, Wolosker H (2005) d-serine is the dominant endogenous coagonist for NMDA receptor neurotoxicity in organotypic hippocampal slices. J Neurosci 25:9413–9417.

Song X, Jensen MO, Jogini V, Stein RA, Lee CH, McHaourab HS, Shaw DE, Gouaux E (2018) Mechanism of NMDA receptor channel block by MK-801 and memantine. Nature 556:515–519.

Stein IS, Gray JA, Zito K (2015) Non-Ionotropic NMDA Receptor Signaling Drives Activity-Induced Dendritic Spine Shrinkage. J Neurosci 35:12303–12308.

Stein IS, Park DK, Claiborne N, Zito K (2021) Non-ionotropic NMDA receptor signaling gates bidirectional structural plasticity of dendritic spines. Cell Rep 34:108664.

Stein IS, Park DK, Flores JC, Jahncke JN, Zito K (2020) Molecular Mechanisms of Non-ionotropic NMDA Receptor Signaling in Dendritic Spine Shrinkage. J Neurosci 40:3741–3750.

Thomazeau A, Bosch M, Essayan-Perez S, Barnes SA, De Jesus-Cortes H, Bear MF (2021) Dissociation of functional and structural plasticity of dendritic spines during NMDAR and mGluR-dependent long-term synaptic depression in wild-type and fragile X model mice. Mol Psychiatry 26:4652–4669.

Ultanir SK, Kim JE, Hall BJ, Deerinck T, Ellisman M, Ghosh A (2007) Regulation of spine morphology and spine density by NMDA receptor signaling in vivo. Proc Natl Acad Sci U S A 104:19553–19558.

Valbuena S, Lerma J (2016) Non-canonical Signaling, the Hidden Life of Ligand-Gated Ion Channels. Neuron 92:316–329.

Wong JM, Gray JA (2018) Long-Term Depression Is Independent of GluN2 Subunit Composition. J Neurosci 38:4462–4470.

Woods GF, Oh WC, Boudewyn LC, Mikula SK, Zito K (2011) Loss of PSD-95 enrichment is not a prerequisite for spine retraction. J Neurosci 31:12129–12138.

Xiao MY, Wasling P, Hanse E, Gustafsson B (2004) Creation of AMPA-silent synapses in the neonatal hippocampus. Nat Neurosci 7:236–243.

Yamada Y, Chochi Y, Takamiya K, Sobue K, Inui M (1999) Modulation of the channel activity of the epsilon2/zeta1-subtype N-methyl D-aspartate receptor by PSD-95. J Biol Chem 274:6647–6652.

Yasuda R (2006) Imaging spatiotemporal dynamics of neuronal signaling using fluorescence resonance energy transfer and fluorescence lifetime imaging microscopy. Curr Opin Neurobiol 16:551–561.

Zhang S, Ehlers MD, Bernhardt JP, Su CT, Huganir RL (1998) Calmodulin mediates calcium-dependent inactivation of N-methyl-D-aspartate receptors. Neuron 21:443–453.

Zheng X, Zhang L, Wang AP, Bennett MV, Zukin RS (1999) Protein kinase C potentiation of N-methyl-D-aspartate receptor activity is not mediated by phosphorylation of N-methyl-D-aspartate receptor subunits. Proc Natl Acad Sci U S A 96:15262–15267.

